# Live-Cell Imaging in Human Colonic Monolayers Reveals Erk Waves Limit the Stem Cell Compartment to Maintain Epithelial Homeostasis

**DOI:** 10.1101/2022.02.23.481374

**Authors:** Kelvin W Pond, Olga Alkhimenok, Jayati Chakrabarti, Yana Zavros, Curtis A Thorne, Andrew L Paek

## Abstract

The establishment and maintenance of different cellular compartments in tissues is a universal requirement across all metazoans. Maintaining the correct ratio of cell types in time and space allows tissues to form patterned compartments and perform complex functions. Patterning is especially evident in the human colon, where tissue homeostasis is maintained by stem cells in crypt structures that balance proliferation and differentiation. Here we developed a 2D patient derived organoid (PDO) screening platform to study tissue patterning and kinase pathway dynamics in single cells across hundreds of conditions. Using this system, we discovered that waves of Erk signaling induced by apoptotic cells play a critical role in maintaining tissue patterning and homeostasis. If Erk is activated acutely across all cells instead of in wavelike patterns, then tissue patterning, and stem cell maintenance are lost. Conversely, if Erk activity is inhibited, then stem cells become unrestricted and expand dramatically. This work demonstrates that the colonic epithelium requires coordinated Erk signaling dynamics to maintain patterning and tissue homeostasis. Our work reveals how Erk can antagonize stem cells yet support cell replacement and the function of the gut.

## Introduction

The colonic epithelium is a highly organized and rapidly renewing tissue containing spatially separated compartments called crypts. Roughly 10 million evenly spaced crypts inhabit the colon and are renewed every 3-5 days [1]. Each colonic crypt is a ~2000 single cell monolayer invagination, roughly 82 cells in height and 41 cells in diameter [2]. Cell extrusion and death are largely confined to non-crypt compartments of the colon, while cell division, required to replace differentiated cell types is restricted to the crypt base. The distance separating these compartments is ~300 microns [3, 4]. Remarkably, chemical destruction of these millions of repeated structures can be repaired in the normal gut within weeks [5], thereby reestablishing spatial separation between stem and differentiated cell compartments with remarkable accuracy. Across these cell compartments, the Wnt and MAPK pathways oppose one another to maintain homeostasis by balancing differentiation and stemness [6–10]. Erk activity is low in colonic crypts and increases as cells become more differentiated. Wnt inhibition leads to hyperactive Erk signaling [8], suggesting that Wnt is suppressing Erk signaling as it maintains stemness. Conversely, loss of Erk1/2 triggers the expansion of stem cells [7], suggesting Erk activity limits the stem cell pool. While much is known about the pathways required for maintaining the different cell types of the colon, how these compartments communicate with each other and maintain regular spacing is unknown.

MAPK/Erk signaling dynamics in wound healing have recently been described. Elegant work in MDCK cells found that cells at the wound barrier initiate Erk signaling cascades in wavelike patterns, which spread away from the site of damage. These Erk waves drive collective movement in the direction of the wound to efficiently seal barriers [11]. Erk activity waves during wound healing have been observed *in vitro* and *in vivo* in a diverse set of animal models, suggesting a highly conserved function in the wound healing process. Erk signaling waves are induced in response to apoptosis [12] and promote hypertrophy [13]. Importantly, Erk dynamics can instruct fate decisions. In neuronal cells and Drosophila embryos, Erk signaling promotes differentiation if the signal is prolonged or total strength is increased [14, 15]. A recent study using organoid monolayers derived from murine small intestine showed that differentiated villus cells have higher Erk signaling over time compared to crypt cells [16]. In summary, Erk signaling can propagate across the epithelium in wave-like patterns in response to cues including apoptosis, regenerative need, and oncogenic mutations [12, 13, 17, 18]. Can Erk waves driven by apoptosis aid the gut in maintaining the proper ratios of cell compartments?

Herein, using self-organizing primary human colon organoid monolayers, we show that epithelial cells can reorganize from single cells into a complex monolayer resembling the tissue of origin. Regularly spaced crypt-like structures (hereafter referred to as “nodes”) form within two days and are maintained in spatially distinct compartments. In non-node compartments, cell extrusion and apoptosis are increased. As a cell is extruded and undergoes apoptosis, an Erk activity wave is initiated that propagates radially from the dying cell. Locally, cells in the path of the wave are prompted to migrate toward the dying cell, presumably to maintain epithelial integrity as reported recently [12, 19]. The spacing of nodes is similar to the diameter of Erk waves, which suggests that the length scale observed between nodes is dependent on Erk waves. Consistent with this, we found that if Erk activity is globally increased, nodes disappear and cease to express the Wnt signaling and stem cell markers LGR5 and MYC. Alternatively, if Erk is inhibited globally, nodes increase in size and express high levels of LGR5 and MYC. If Wnt3a is removed from culture media, nodes also disappear and Erk activity is increased. These data show that Wnt and Erk mutually oppose one another to maintain gut homeostasis and Wnt-suppressing Erk waves originating from apoptotic cells help to shape the architecture of the human colonic epithelium.

## Methods

### Preparation of 3D organoids from Patient Biopsies

Normal or tumor tissues from Endoscopic ultrasound-guided fine-needle aspiration biopsies (EUS FNAs), or core needle biopsies were collected from consented patients by the TARGHETS (Tissue Acquisition and Repository for Gastrointestinal and HEpaTic Systems, IRB 1909985869) facility located in the Arizona Health Science Center. Biopsy tissues are transported in 50ml conical tubes containing collection media (Advanced DMEM/F-12 (Invitrogen 12634028) supplemented with 2 mM GlutaMax, 10 mM HEPES, 0.25mg/ml Amphotericin B, 10mg/ml Gentamycin, 1% Kanamycin, N-2 media supplement (Invitrogen 17502048), B-27 Supplement Minus Vitamin A (Invitrogen 12587010), 1mM N-Acetyl-L-cysteine (Sigma A9165), 10nM Nicotinamide (Sigma Aldrich; #N0636), 2.5µM CHIR99021 (Tocris-Fisher, 4423; Apexbio Technology, A3011), and 2.5 µM Thiazovivin. Tissues were then washed, minced, and cryopreserved in organoid freezing media (70% seeding media (see next section), supplemented with 20%FBS, 10% DMSO, and 2.5 µM Thiazovivin) at the BioDROids core facility, located in the University of Arizona Cancer Center. Frozen tissues were prepared into organoids by thawing tissue pieces, mincing, and incubation with 1mg/ml Collagenase Type 3 in PBS on a shaker for 10-25 minutes at room temperature. Cells were then removed from collagenase, washed with PBS, embedded into 100% Matrigel, and cultured in seeding media for 7 days. After organoids became established, low passage aliquots were cryopreserved in freezing media or infected with lentiviral constructs.

### Growth Conditions of 3D and organoid monolayers

#### Complete LWRN media

Advanced DMEM/F-12 (Invitrogen 12634028) supplemented with 2 mM GlutaMax, 10 mM HEPES, N-2 media supplement (Invitrogen 17502048), B-27 Supplement Minus Vitamin A (Invitrogen 12587010), 1mM N-Acetyl-L-cysteine (Sigma A9165), 2500 units/mL Penicillin and 2.5 mg/mL streptomycin (0.05 mg/mL), 50% L-WRN Conditioned Media [3], 100 ng/ml huEGF (R&D 236-EG-01M), 500nM A 83-01 (Tocris-Fisher, 29-391-0; APexBio-Fisher, 501150476), 10µM SB 202190 (Tocris-Fisher, 12-641-0),100µg/ml Primocin, and 10mM Nicotinamide (Sigma Aldrich; #N0636).

#### Seeding media

Complete LWRN media supplemented fresh with 2.5µM CHIR99021 (Tocris-Fisher, 4423; Apexbio Technology, A3011) and 10µM Y27632 (Tocris-Fisher; 125410).

### Lentiviral Infection of 3D Organoids

3×10^6^ HEK 293T cells were seeded in a 10cm plate and transfected the following day using 500μL of opti-MEM, 30μL Gene Juice (Sigma #70967), 5ug lentiviral construct, 3.25µg psPAX2 (addgene #12260), and 1.75µg pMD2.G (addgene #12259). After 24 hours, fluorescence was assessed, and media was collected for 3 days followed by centrifugation and filtration using 0.45µM syringe filter. Lentiviral media was then kept at 4°C before 100x concentration using LentiX concentrator (Takara #631232) and resuspension of virus in organoid seeding media, viral media was then stored at −80deg C for up to 3 months. For infection of organoids, 1×10^5^ primary epithelial cells were harvested from 3D domes, washed, and trypsinzed as described previously. Cells were then counted using a hemocytometer and 40K organoids were plated in suspension onto a 48 well plate and diluted 1:1 in vial media containing 8mg/ml polybrene. Next, organoids were placed at 37°C for 1hour the before centrifugation at 600g for 1hr at 32°C. Organoids were then harvested, washed, and embedded into 90% Matrigel and cultured in seeding media for 24 hours. Cells were then grown in complete LWRN for 3 days before fluorescence was assessed and >50% infection was achieved. Aspirate media and re-suspend organoids in 500ul Trypsin. After dissociation, cells we mixed 5-10 times using a p200 pipette tip and passed through a 100μm filter to ensure isolation of single cells.

### Preparation of organoid monolayers from 3D Cultures

SCREENSTAR 384-well black plates (Grenier #781866) were coated with ice cold Matrigel^™^ (CB40230C) diluted 1:40 in Serum-free Advanced-DMEM F12 media (SFM) for 1 hour. Monolayers were prepared using our previously described protocol [20, 21]. Briefly, cells were removed from 3D Matrigel and resuspended in ice cold SFM and washed three times using ice cold SFM. Organoids were then dissociated by resuspension in Trypsin+10 µM Y-27 for 4 minutes followed by quenching in DMEM supplemented with 10% FBS and washing. Coating media was removed from 384-well plates and organoids plated @ 7000 cells/well after counting using hemocytometer and cultured for 24 hours. Seeding media was then removed and replaced with complete LWRN media and changed daily for 7 days until cultures became confluent and tissue patterning was observed.

### High Content Imaging of organoid monolayers

384-well plates were imaged with fluorescent microscopy on a Nikon Eclipse TI2 automated microscope or. For quantification of wave dynamics, organoids were imaged every 10-30 minutes. Tracking and segmentation of single cells in time lapse images in Figure 2 was performed using the MATLAB program p53Cinema [22]. Analysis of the extracted data was performed in MATLAB. For the Erk activity heat maps in Figure 3, images were captured every 7 minutes for 24 hours. For each frame, individual nuclei were segmented using a MATLAB script developed in-house. Cytoplasmic regions were defined by a donut-shaped ring two pixels wide surrounding the nucleus. Cells with active ERK were identified as having a mean ERK-KTR cytoplasmic signal => the ERK-KTR nuclear signal. The density-based spatial clustering of applications with noise algorithm (dbscan, MATLAB, 2021b) was performed on the centroids of ERK active cells to identify spatial clusters of cells with active Erk. A boundary was drawn around the spatial clusters using the boundary algorithm in MATLAB. The Erk positive regions of each frame were summed up to generate the heat map in MATLAB.

### Plasmids

1. pLentiPGK DEST H2B-iRFP670 was a gift from Markus Covert, addgene #90237 [4].
2. pLentiPGK BLASTDEST ErkKTRmRuby2 was a gift from Markus Covert, addgene #90231 [4].
3. PGF-H2BeGFP was a gift from Mark Mercola, addgene #21210 [23].

#### Antibodies and Dyes

Goat polyclonal anti-Muc2 (Santa Cruz Biotechnology, sc-23170)

Alexa Fluor 546 Phalloidin (Thermofisher, A12379)

Click-iT™ EdU Cell Proliferation Kit for Imaging, Alexa Fluor™ 488 dye (Thermofisher, C10337)

DAPI (Thermofisher, D21490)

Rat monoclonal anti-Ki67 (Thermofisher, 14-5698-80)

Goat anti-Rat IgG (H+L) Cross-Adsorbed Secondary Antibody, Alexa Fluor 488 (Thermofisher, A-11006)

Donkey anti-Goat IgG (H+L) Secondary Antibody [DyLight 488] (Pre-adsorbed) (Novus, NBP1-74824)

NucView® Caspase-3 Enzyme (Biotium #10403)

#### Fluorescent-assisted Cell Sorting (FACS)

After infection with lentiviral reporters, dual reporter organoids were derived. Organoids were dissociated into single cells and sorted for similar levels of expression for the desired reporter (see figure S1C for representation of FACS plots generated using FloJo V10). Cell sorting was performed using the BD FACS Aria III flow cytometer.

#### scRNA FISH

Experiments were done using previously validated and custom RNA probes for target mRNA of interest using Molecular Instruments HCR RNA FISH protocol for mammalian cells [24] (Choi, Nat Biotech, 2010). Briefly, monolayers were fixed in 4% PFA, permeabilized in 70% EtOH at −20degC, hybridized at 37deg C, and amplified overnight at RT. Monolayers were then imaged using a Nikon CSU-W1-Sora Spinning Disk Confocal Microscope. See Supplementary Table 1 for complete probe sequences.

## Results

### Patient-derived Colonic Organoid Monolayers

To study single-cell dynamics in the human colon, we adapted our previous method of culturing murine intestinal organoids as monolayers. These monolayers maintain patterning and cell fate characteristics observed in 3D organoid systems and in vivo [20]. In conjunction with the University of Arizona Cancer Center organoid resource (BioDROid), we collected colonic normal, adenomatous polyp, and carcinoma samples. We cultured them as 3D patient-derived organoids (PDOs), expanded them in 3D and transferred them onto a thin layer of extracellular matrix after dissociation into single cells. After four days, monolayers were considered fully established (Figure 1A). Strikingly, even starting as a random dispersion of single cells, normal colonic organoids showed localized compartments of densely packed cells (nodes) that were surrounded by less dense cells. These nodes resembled those we observed using the murine model system [20], with LGR5+ stem cells located specifically within densely compacted cell compartments (figure S1). The murine small intestinal organoid monolayers as well as the normal human colon monolayers rarely reach confluence and instead reach a homeostatic state and maintain nodes at sub-confluent levels (Figure S2). A polyp-derived 2D organoid line (referred here as GiLA1 for <u>G</u>astrointestinal <u>L</u>ine, <u>A</u>rizona <u>1</u>) formed regularly spaced nodes, quickly reached confluence within 72 hours, and could maintain node structures for 45 days or more. When GiLA1 single cells were seeded onto an ultra-thin layer of Matrigel, organoids formed monolayers by three days and began to spontaneously form regularly spaced compact compartments reminiscent of colonic crypts within 4-5 days (Figure 1C and 1D). A tumor organoid derived from an MSI-negative invasive colon adenocarcinoma (p21T) was grown in 2D and displayed different morphologies compared to normal, polyp, and the murine small intestine (Figure S2). p21T proliferated but failed to form nodes and instead formed uniform disorganized monolayers (Figure S2). To date, across multiple patient-derived organoid lines, we have observed the formation of nodes in normal and polyp-derived organoids; however, we fail to observe nodes in multiple adenocarcinoma organoids.

**Figure 1.**
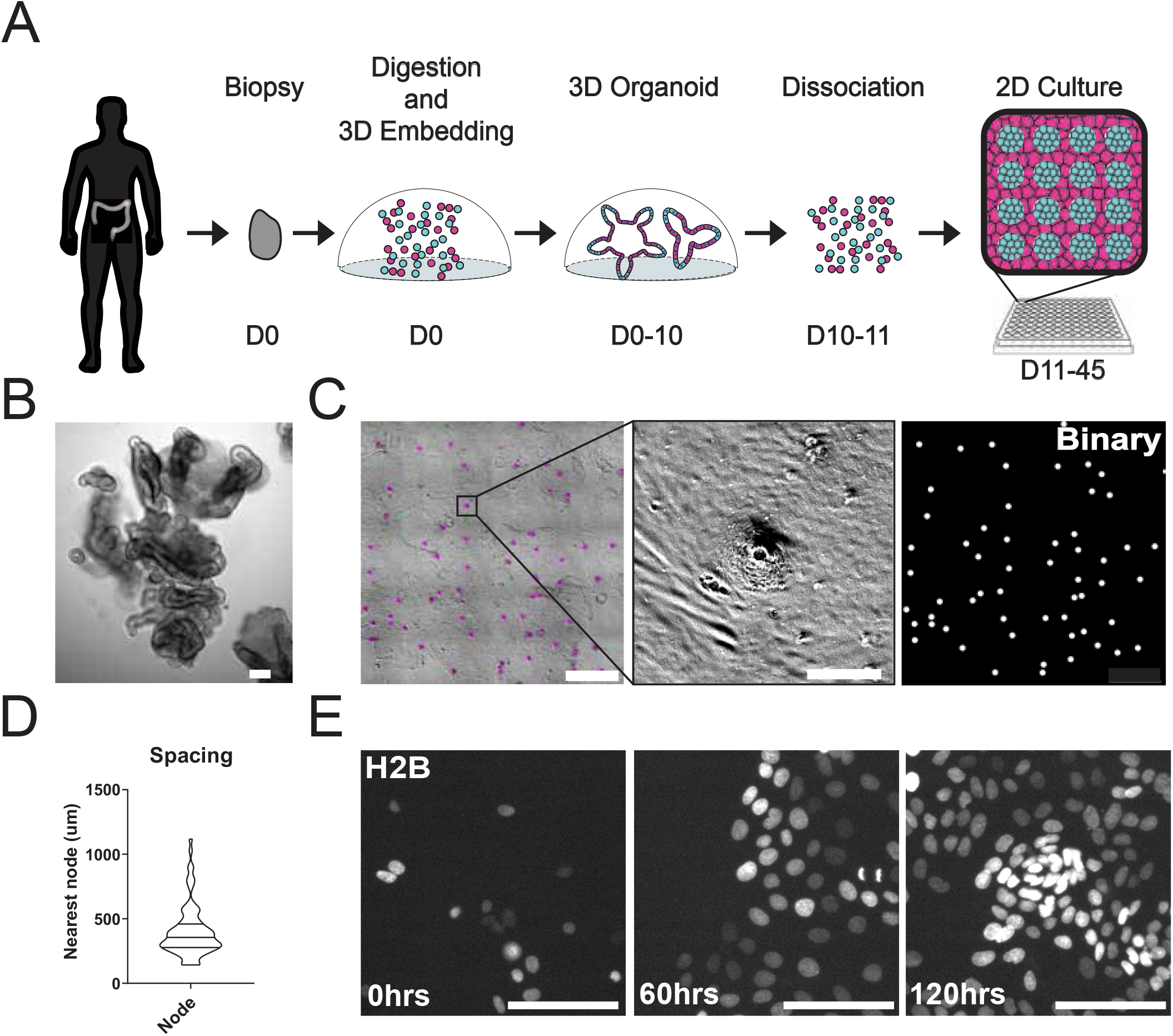
Development and Self-organization of Colonic Patient-derived Colonic Organoid Monolayers. A. Model depicting the workflow for the development of organoid monolayers. Patient biopsies are digested using collagenase to form a single cell suspension. Cells are then embedded in Matrigel and grown in 3D using a defined organoid growth medium. Organoids are expanded in 3D before seeding onto 2D imaging plates coated with a thin layer of Matrigel. Within 5 days, single cells self-organize to form a regularly patterned organoid monolayer. Organoid monolayers maintain homeostasis in this patterned state for up to 45 days. B. Representative brightfield image of a single 3D colonic organoid. C. Representative brightfield image of organoid monolayer. Left-automated image segmentation of nodes across a single well. Automated detection of nodes is shown in purple. Middle-Single node, zoom of left. Right-Binary image of segmentation showing regular spacing of nodes across the culture. D. Violin plot showing the distribution of spacing across organoid monolayers. E. Live cell images of self-organization over 5 days using organoids expressing H2B-iRFP670 for nuclear tracking. All scale bars represent 100µM except C, left, which represents 1000µM.

### Self-organization of Organoid Monolayers

A striking aspect of the organoid monolayers is that they appear to self-organize into node/non-node compartments. To capture the establishment of these structures over time, we used time-lapse microscopy of GiLA1 organoid monolayers expressing H2B-iRFP670 to observe the formation of distinct cellular compartments over five days. Cells initially exhibited high levels of motility after attachment to the extracellular matrix (ECM) to form 5-10 cell clusters. After this, cells became hyperproliferative from 48-72 hours after seeding to form a complete monolayer. Finally, morphologically distinct cellular compartments were formed after 96 hours (Figure 1E, Movie 1). These 2D monolayers can be readily grown in 384-well imaging plates making this model particularly amenable to high content imaging and quantitative image analysis. Together, these data show that patient biopsies from the human colon can fully self-organize into patterned monolayers that maintain long-term tissue homeostasis.

### Spatially Distinct Stem and Differentiated Cell Compartments

To study the cell types the node structures harbor and how they form, we further characterized the GiLA1 organoid line. Two proliferative markers, Ki-67 and EdU, revealed that proliferative cells were much more abundant within the compartments of densely packed cells of the organoid monolayers (Figure 2A and 2C), suggesting the nodes are a collection of transit amplifying-like cells and possibly stem cells. To determine if nodes harbored stem cells, we performed single cell RNA fluorescence in-situ hybridization (scRNA FISH) against Wnt target genes and stem cell markers, LGR5 and MYC. LGR5 and MYC expressed at higher levels in nodes compared to non-nodes (Figure 2B and 2D). Cells expressing the differentiated-colonocyte marker KRT20 were primarily located in non-node regions and were mutually exclusive to LGR5 high compartments (Figure 2B and 2D). These data show that patterned organoid monolayers harbor cells positive for either stem, proliferative, or differentiated cell markers.

**Figure 2.**
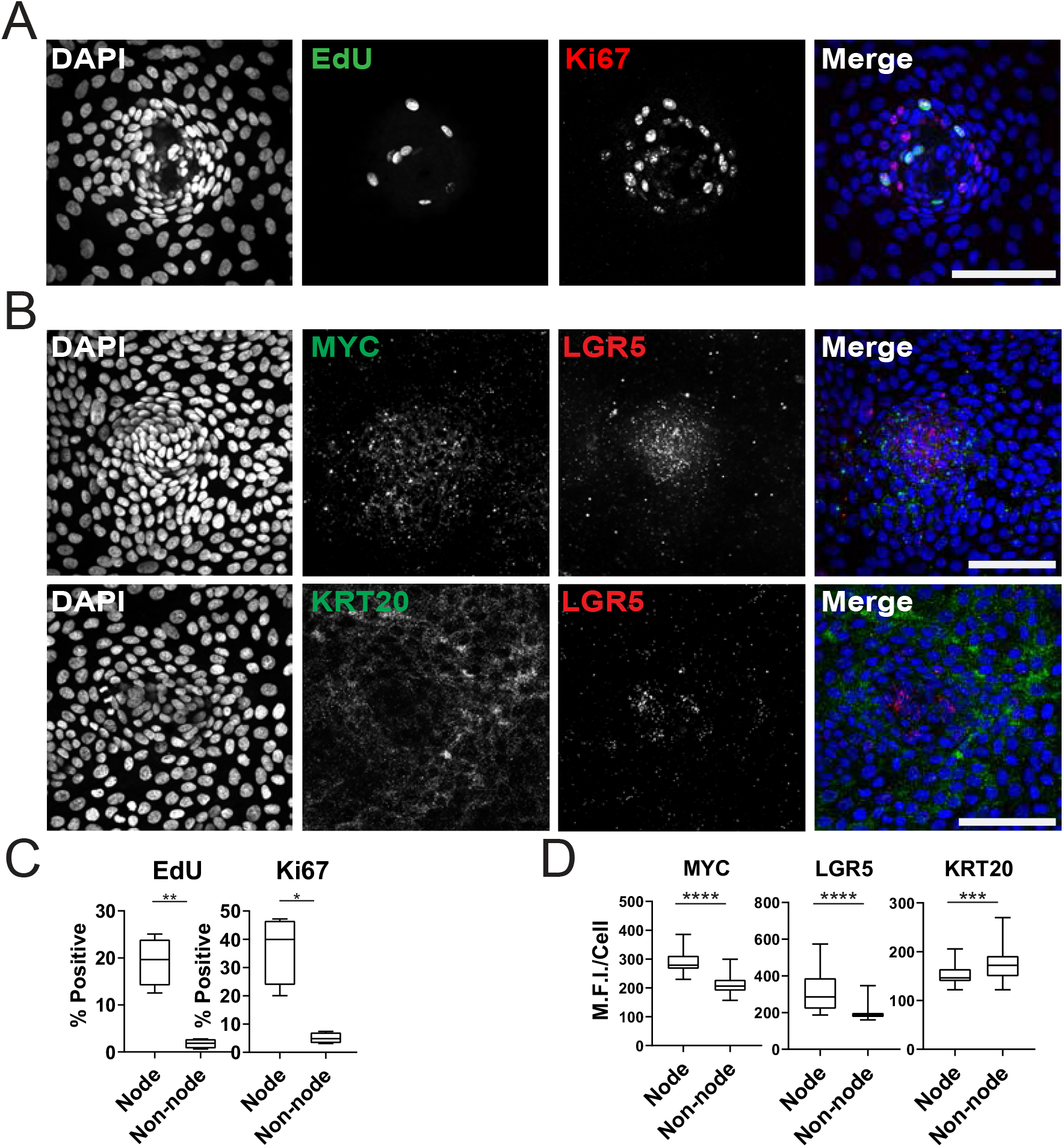
Characterization of Organoid Monolayer Compartments. A. Representative images 2D organoids stained for nuclei (blue), EdU (green), and ki67 (red). B. Representative images of single-cell RNA FISH against Wnt target genes MYC and LGR5 in a single node. C. Quantification of A showing % positive cells in node or non-node regions, asterisks represent significance from paired t-test. D. Quantification of images represented in B, mean fluorescence intensity (MFI) of target transcripts in each single cell is shown. Cells were separated based on presence in nodes vs non-nodes. Asterisks represent significance from Mann Whitney analysis. All scale bars represent 100µM.

### Apoptosis-induced Erk Waves Instruct Cell Movement

MAPK/Erk signaling is a critical regulatory pathway involved in maintaining homeostasis of epithelia in the gut [7]. To assess Erk activity in human organoid monolayers over time, we utilized a well-characterized Erk kinase translocation reporter (ErkKTR) [25]. The ErkKTR utilizes a bipartite NLS and NES to transform Erk kinase activity into a nuclear to cytoplasmic shuttling event that can be easily quantified by microscopy in organoid monolayers (Figure 3A). We transduced the GiLA1 organoid line with a H2B-iRFP670 nuclear marker and ErkKTR-ruby2 reporter to perform nuclear and cytoplasmic segmentation and calculate nuclear-to-cytoplasmic ratios of the ErkKTR (Figure 3A, Figure 3B and S3). In fully developed organoid monolayers, we observed a ~2-fold reduction of the ErkKTR after treatment with inhibitors of EGFR or MEK across all cell types, suggesting the KTR is actively representing Erk signaling and is EGFR dependent (Figure S4). Interestingly, when GiLA1 monolayers were imaged using time lapse microscopy, we observed Erk activation originating at focal points within the confluent monolayer that propagate outward in a wave-like pattern. These waves did not overlap and were observed routinely during homeostasis (Figure 3C, Movie 2). Imaging of monolayers using brightfield and caspase 3 dye revealed apoptotic cells that were being extruded from the monolayer to be the focal source of wave activation (Movie 2).

**Figure 3.**
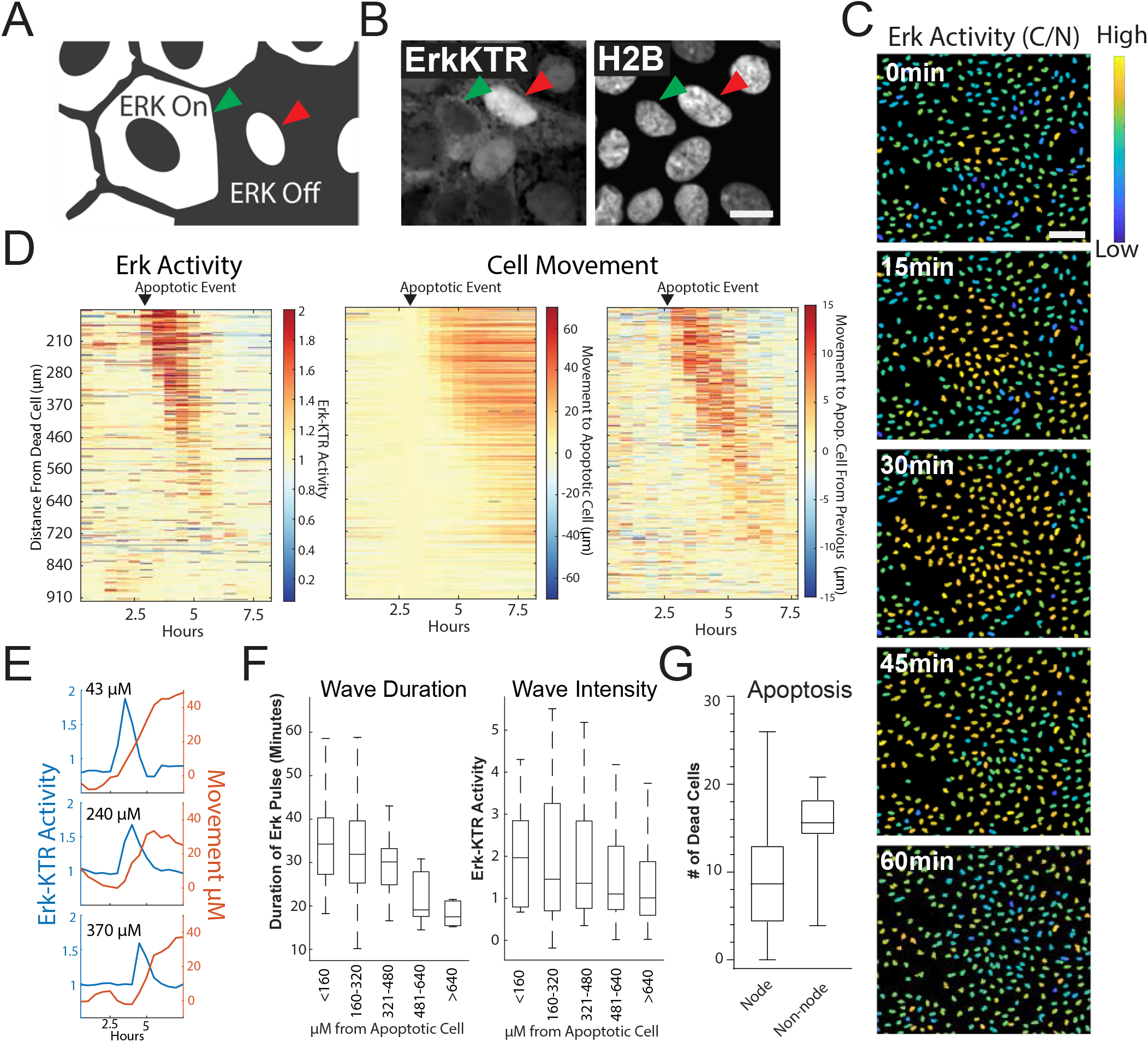
Apoptosis Induces an Erk Signaling Wave that Instructs Cell Movement. A. Model depicting how to interpret the ErkKTR translocation reporter. Cells with mostly cytoplasmic ErkKTR have high Erk activity (Left, green arrow). Cells with mostly nuclear ErkKTR signal have low Erk activity (Right, red arrow). Scale bar represents 10µM. B. Representative images of the ErkKTR active versus inactive. C. Representative images showing a single Erk Wave propagating from an apoptotic cell. Nuclear to cytoplasmic ratio ErkKTR heat map is shown in blue (low)/yellow (High). Scale bar represents 100µM. D. Left-single cell analysis of Erk activity over time after an apoptotic event. Heat maps are ordered from closest (top) to furthest (bottom) distance from the dying cell over time. Erk activity is shown (Red, high Erk activity; Blue, Low Erk activity). Middle-single cell analysis of cell movement obtained from the same dataset shown on left. Change in distance towards the position of the apoptotic event is shown. Right-relative change in distance compared to the previous frame is shown. E. Representative single cell traces comparing Erk activity (blue) and cell movement (red) at a given distance from the apoptotic cell is shown. F. Left-duration of Erk signaling wave at a given distance from an apoptotic cell. Right-ErkKTR activity at a given distance from an apoptotic cell. G. Location of cleaved caspase 3 positive cells in monolayer over 24 hours. Data represented as the number of caspase positive cells in node or non-node area.

To determine the properties of the Erk activity waves, we tracked ~450 cells and assessed their Erk activity over time following an apoptotic event (Figure 3D, left panel, 3E). Both apoptotic events initiated a wave of Erk activity that traveled approximately 450 µM over a period of 95 minutes for an average speed of ~4.7 µM/minute. In addition to the Erk signaling wave, we noticed apparent cell movements at the site of apoptosis. Tracking single cells revealed a striking correlation between intensity/duration of Erk activity and movement of cells surrounding the apoptotic cell in the direction of the apoptotic cell (Figure 3D, two right panels and 3E). As the Erk wave propagated outward from apoptotic cells, it dissipated in both duration and intensity (Figure 3F). Particle image velocimetry (PIV), a method to quantify image movement over time, confirmed collective cell migration towards the apoptotic cell (Figure S5), suggesting that Erk waves instruct migration of surrounding cells as shown in previous studies [12]. Cells treated with the EGFR inhibitor Gefitinib did not induce Erk signaling waves or display tropism towards dying cells, suggesting that EGFR-mediated Erk signaling is required for cell movement in this context (Figure S5, Movie 3). A cross correlation analysis revealed that Erk activity preceded cell movement (Figure S6), suggesting that Erk waves instruct cell movement. We previously described distinct apoptotic and proliferative compartments in organoid monolayers derived from murine small intestine [20]. To confirm this in human organoid monolayers and investigate if cell death was restricted to a compartment distinct from proliferative nodes, we imaged mature living organoid monolayers in the presence of cleaved caspase-3 dye, a marker for apoptosis. We found that the emergence of an apoptotic cell was more likely in non-node compartments, largely restricted to the differentiated cells (Figure 3G and S7). Taken together these data show that Erk waves induced by apoptotic cells instruct surrounding cells to migrate toward the apoptotic site.

### Erk waves instruct patterning in organoid monolayers

Increased Erk activity has been reported in differentiated cells compared to the stem cell compartment within the gut [8]. We asked if Erk activity was increased in the differentiated compartment in GiLA1 monolayers. To determine this, we created a time lapse movie of a dense node compartment surrounded by less dense, non-node cells over a 16-hour period. Nuclear to cytoplasmic ratio of the ErkKTR was measured at the single cell level over time. A threshold for active Erk cells was then combined with spatial clustering to identify Erk-active regions. High Erk regions were then superimposed from each imaging timepoint to create a single heat map for regional Erk activity over time. Consistent with prior studies using murine small intestine [8], cells in non-node compartments were more mobile and displayed more active Erk signaling (Figure 4A and Movie 4). Together, these data show that apoptotic events that cause Erk activity waves are mostly localized in spatially distinct differentiated cell compartments.

**Figure 4.**
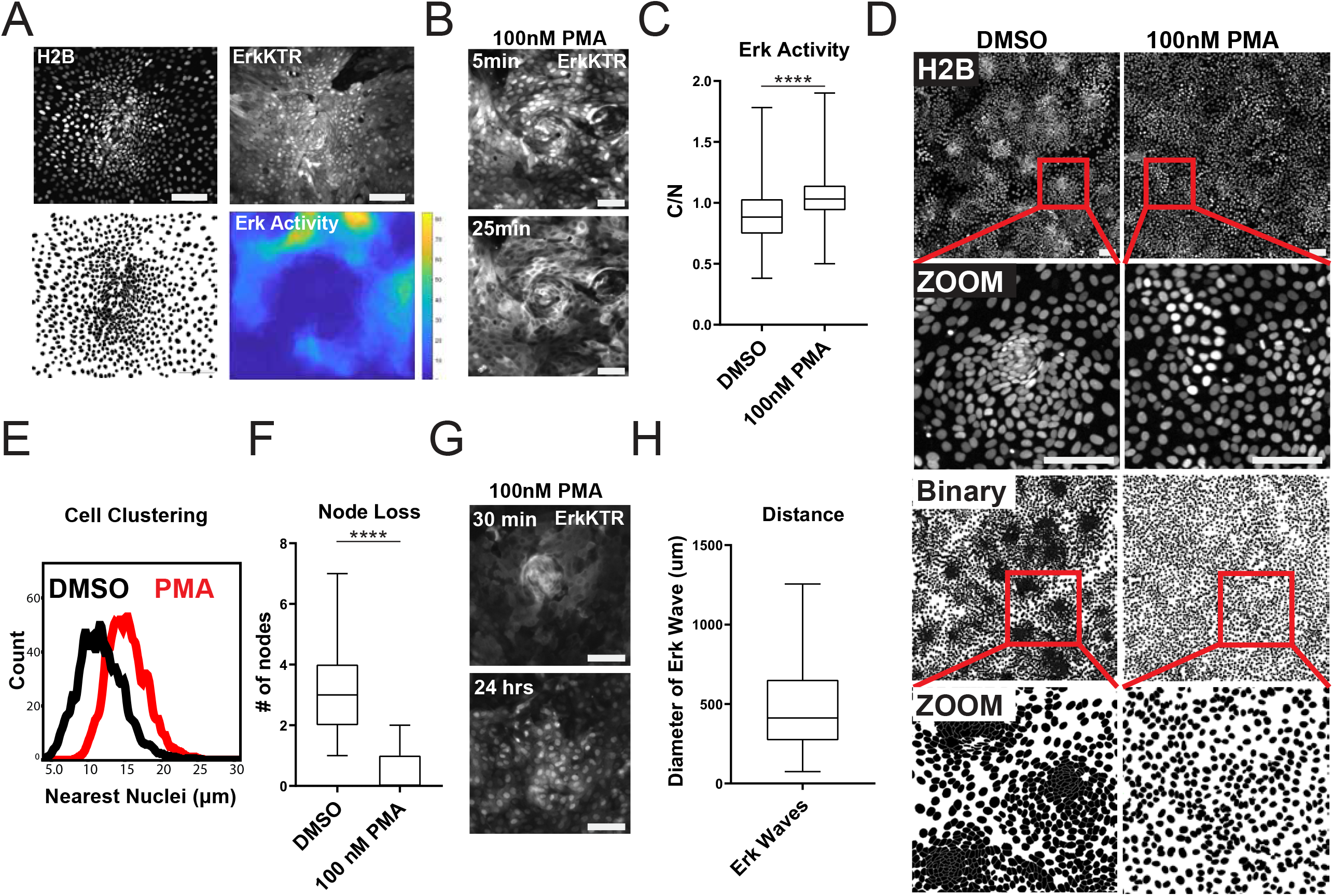
Erk Dynamics are Essential to Maintain Tissue Patterning in Organoid Monolayers. A. Representative images of cell density variation in node versus non-node. Top left-H2B-iRFP670. Top right-ErkKTR. Bottom left-binary nuclear segmentation showing high- and low-density regions. Bottom right-a spatial heat map of Erk activity over 16 hours in node and surrounding region. B. Representative images of Erk activation after 100nM PMA treatment for 25 minutes. Top-ErkKTR 5 minutes after treatment with PMA. Bottom-ErkKTR of the same node 25 minutes after PMA treatment. C. Quantification of ErkKTR activity after 30 minutes of 100nM PMA treatment. D. Cell clustering following 100nM PMA treatment. Top 4 panels show nuclear distribution across stitched image (top) and zoom (bottom) before and after PMA treatment. Bottom 4 panels show binary segmentation to clearly show loss of cell clustering following PMA treatment. E. Histograms showing distance between nuclei with or without treatment with 100nM PMA. Data is represented as nearest nuclei distance. F. Quantification of patterning loss before and after treatment with 100nM PMA for 24 hours. G. Representative images of ErkKTR activity after pulsing cells for 1 minute with PMA followed by washout and chase for 24 hours. Top-all cells show high Erk activity with intact node. Bottom-node loss and re-suppression of Erk activity after 24 hours. H. Quantification of distribution of Erk wave diameter in organoid monolayers. Asterisks represent significance from Mann Whitney analysis. Scale bars represent 100µM.

Since apoptotic events and Erk activity are largely restricted to the post-mitotic, differentiated compartments in our gut organoid monolayer model, we hypothesized that Erk waves may help regulate the patterning observed in the GiLA1 monolayers by restricting the stem cell compartment. To test if Erk dynamics help to maintain the patterning of the organoid monolayers, we asked if global activation of Erk activity could disrupt the patterning of the organoid monolayers. We used a well-established activator of the Erk pathway, Phorbol 12-myristate 13-acetate (PMA) to acutely activate Erk signaling across the monolayer. Following treatment with 100nM PMA, we observed immediate hyperactivated Erk in all cells (Figure 4B and 4C). Monolayers showed a remarkable loss of patterning after PMA treatment as evidenced by almost complete loss of cell clusters by 24 hours post treatment (Figure 4D-4F). A similar experiment where monolayers were pulsed with 100nM PMA for 1 minute revealed Erk activation in all cells after ~30 minutes, also to trigger node loss. Notably, Erk levels returned to normal after PMA treatment but nodes did not re-form by 48 hours (Figure 4G and Movie 6). This data suggests a potential role for apoptosis induced Erk signaling in shaping the architecture of the colon. In support of this hypothesis, we observed a striking similarity in the average distance (~233.7 μm) between neighboring proliferative compartments (Figure 1D) and the average distance (~238 μm) covered by the apoptosis-induced Erk wave (Figure 4H).

Wnt is a well-known promoter of stem cell renewal within the gut and may be responsible for the clustering of the proliferative nodes as we previously observed in murine small intestine monolayers [20]. We asked if removal of Wnt from growth conditions would affect stem cell numbers from our GILA1 monolayers. Using scRNA FISH towards stem cell marker LGR5, we observed node structures were enriched in LGR5+ cells and these cells were reduced if Wnt3a was removed for 4 days from organoid culture media (Figure 5A and 5B). Erk activity was also significantly increased as cells became more differentiated following Wnt removal (Figure 5C and 5D, Movie 7), consistent with previous work showing increased Erk within crypts following Wnt inhibition [8]. To measure the direct effects of either inhibition or activation of the Erk pathway across all cell types, we treated the GILA1 monolayers with either a MEK inhibitor or PMA for 72 hours. Strikingly, blocking the Erk pathway increased the expression of LGR5 and resulted in larger clusters of densely packed cells, suggesting Erk waves limit stem cell expansion (Figure 5E and 5F). Conversely, treatment with PMA resulted in almost a total loss of stem cells and node structures (Figure 5E and 5F). Similar results were observed when measuring MYC transcripts in the same context (Figure S8). Collectively, these data support a model where Erk signaling, induced by cell death, promotes differentiation, and restricts stem cells to a defined compartment. Interrogation of either the Wnt or Erk pathways revealed a mutually antagonistic relationship between these central regulatory pathways which maintains homeostasis.

**Figure 5.**
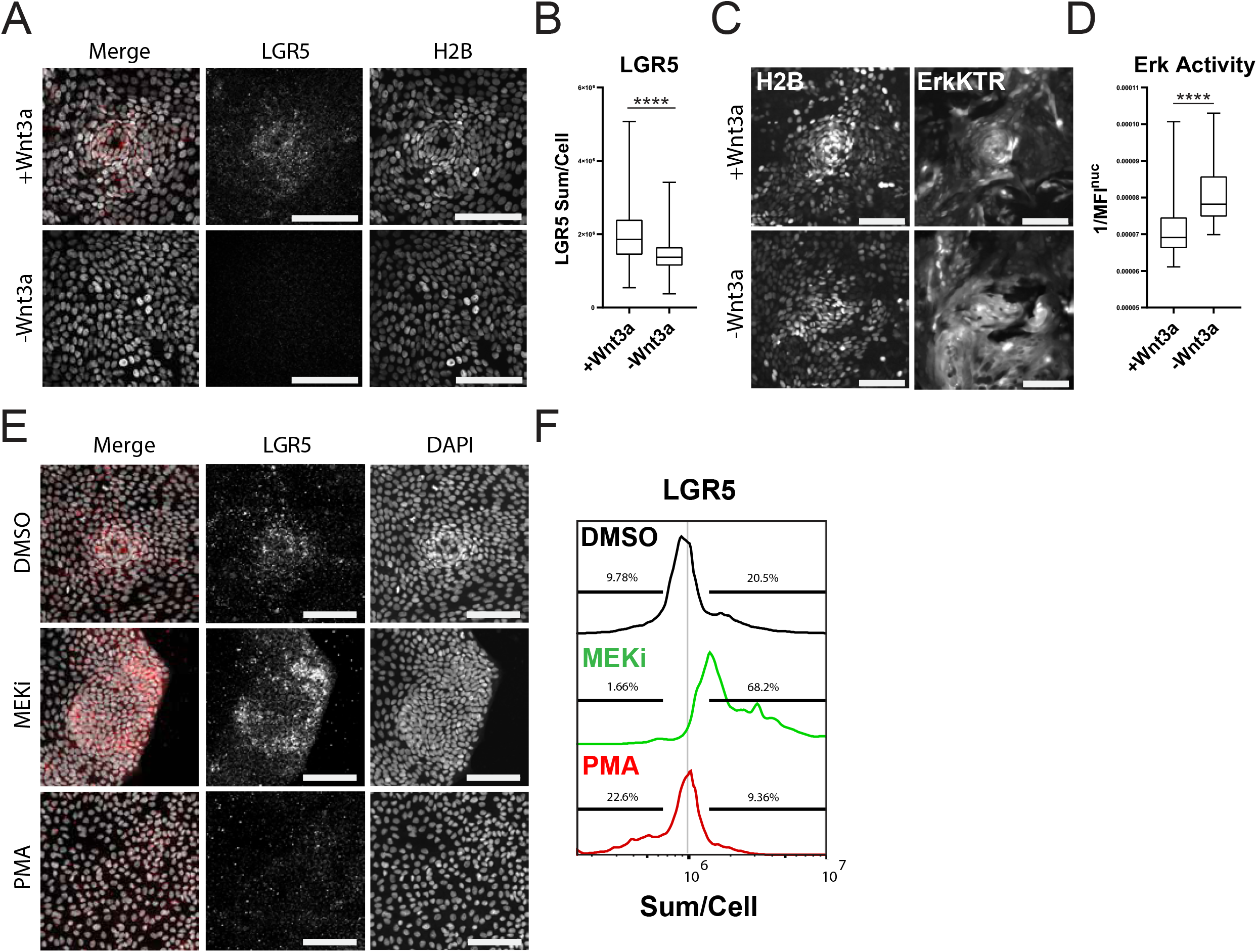
Wnt and Erk Signaling mutually limit each other to preserve tissue homeostasis. A. Representative images of LGR5 mRNA in organoid monolayer after removal of Wnt3a for 72 hours. H2B-GFP is shown in white and LGR5 mRNA is shown in Red. B. Quantification of scRNA FISH for LGR5 in organoid monolayers after removal of Wnt3a from organoid media. C. Representative images of Erk kinase activity after Wnt2a removal for 24 hours. D. Quantification of Erk kinase activity after removal of Wnt3a from organoid media from 0-48 hours following Wnt removal. Inverted nuclear intensity of ErkKTR is shown. E. Representative images of scRNA FISH for LGR5 in organoid monolayers after treatment with either 1µM PD0325901 (MEKi) or 100nM PMA for 24 hours followed by 24 hours of normal media. F. Quantification of experiment described in E. Data is represented as histograms representing the sum of Lrg5 intensity per cell. Gates show percent of high and low LGR5+ expressing cells. Asterisks represent significance from Mann Whitney analysis. Scale bars represent 100µM.

## DISCUSSION

### Erk Waves in the Human Gut

The human colon requires ongoing regeneration and renewal over a lifespan; This need relies on finely tuned communication between differentiated and stem cells. We show here that one way the colonic epithelium maintains both complexity and plasticity is through long distance signaling driven by cell turnover. As cells differentiate and are eliminated from the epithelium, dying cells initiate an Erk wave signal that triggers local cell tropism towards the death event and stimulates differentiation of replacement cells. This mechanism is an efficient way for the colon to maintain the correct proportions of stem and differentiated cells in distinct spatial compartments.

### Organoid Monolayer System

We have shown that primary human colonic tissue can be expanded and transformed into monolayers which phenocopy key characteristics observed in the tissue of origin. Although the immune and stromal compartments are not included in this culture method, different compartments of stem and differentiated cells can be established and form unique physical properties depending on the source of the tissue (Figure 1 and S1B). The role of the stromal and immune compartments in the maintenance of tissue patterning and their effect on Erk dynamics is an exciting area of future study. The mechanism that drives morphological memory from the tissue of origin remains unclear, but we hypothesize dynamic behavior in critical regulatory kinases like Erk play a major role.

### Potential Self-organization Mechanisms

Although our system is self-organizing, this work focuses more on the maintenance of stem and differentiated cell compartments in space. One future step is defining the mechanism by which organoid monolayers can establish these compartments from dissociated single cells. Recent work has highlighted mechanically mediated transcription factors and substrate stiffness as critical mediators of pattern formation in murine small intestine [26–28]. Dissociation of 3D organoids into single cells most likely also triggers a wound healing response resulting in the production of fetal-like wound-associated epithelial (WAE) cells [29–33]. These cells would aid in monolayer formation during self-organization (Figure 1G), then likely disappear as compartments are established. In the future, an air liquid interface (ALI) culture with stromal cells present would limit wound healing and hypoxia phenotypes as described previously [34]. However, the expression of localized Wnt target genes Lgr5 and Myc (Figure 1F), suggests that a fetal-like state has been repressed as the organoid monolayers mature and form patterns [32, 33].

### Apoptosis-induced Survival

Previous work highlighted that apoptosis induced Erk signaling waves limit cell death to maintain epithelial integrity in a process termed apoptosis-induced survival (AiS) [12, 19]. In our organoid monolayer system, dying cells also triggered Erk waves, which directed cell movement toward the site of cell death (Figure 2). Although we did not calculate the probability of cell death in recent wave compartments, we observed a regular spacing of apoptotic cells thus supporting a death refractory zone created by Erk waves.

### Colon architecture

Erk waves have been highlighted recently as central drivers of tissue architecture. The cochlear duct, which relies on its curved tissue architecture to function, is shaped by Erk signaling waves during development [35]. Erk waves are also essential for proper regeneration of zebrafish scales [13]. Similarly, here we show that the spacing of stem cells in colonic organoid monolayers is correlated with the size of differentiation promoting Erk waves. As extruded cells induce these Erk waves, they carve out a differentiated compartment, which drives cells towards apoptosis, resulting in positive feedback. These signals also limit stem cells, providing an elegant mechanism for separating regions of cell death from the stem cell compartment. We find it interesting that Erk waves that occur in zebrafish scale regeneration drive cellular hypertrophy [13]. Similarly, we find that increasing Erk activity with PMA eliminated the stem cell compartment and concordantly caused cellular hypertrophy. It is possible that cellular hypertrophy induced by Erk might cause a reduction in proliferation as an increase in cell size has been causally linked to slower proliferation rates in both yeast and mammalian cells [36, 37]. However, further experiments are needed to confirm this. It would be interesting to determine if Erk waves differ in size based on the tissue of origin.

### Maintaining a balance between stemness and differentiation

In epithelial monolayers, Erk waves are shut off as cells cease to move towards the dying cell through negative mechanical feedback [38]. Here we see that acutely inducing Erk signaling across all cells destroys the stem cell niche, suggesting this mechanical feedback is essential to maintain stem cell homeostasis. Conversely, inhibition of the Erk pathway unleashes stem cells resulting in aberrant expansion of the stem cell compartment. This suggests that the dynamics and location of Erk signals help to maintain the balance between stemness and differentiation. Recent studies in animal models support a mutually inhibitory relationship between the Wnt and Erk pathways in the normal gut. In the murine small intestine, Erk activity is low in the crypts of Lieberkühn and higher as cells move out of the crypt. Inhibition of paracrine Wnt signaling results in Erk hyperactivation in the crypt and loss of LGR5^+^ stem cells [8], suggesting that Wnt is suppressing Erk signaling as it maintains stemness. Inversely, if Erk 1/2 is knocked out at embryonic stages in mice, stem cells within the crypt proliferate and expand dramatically, displaying a ~2-fold increase in LGR5 and OLFM4 [7]. This phenotype is strikingly similar to what we have observed after chemical inhibition of the Erk pathway in our organoid monolayers (Figure 4E). During hair follicle development, a well-established patterning model system, EGFR is essential and serves to limit cell proliferation and stem cell numbers through attenuation of Wnt signaling [6]. β-catenin is also essential for follicle formation and hyperactivation of the Wnt pathway results in de novo follicle expansion [39]. Together these studies support our hypothesis that these two pathways balance one another to maintain tissue homeostasis. Our organoid monolayers lack stromal and immune cells, suggesting that the maintenance of patterning is most likely driven by signaling between epithelial cells rather than stromal or immune cell interactions with the colonic epithelium.

### Implications in Human Disease

The Wnt and Erk pathways are commonly hyperactivated during the development of colon cancer. This work unveils a relationship between these two pathways that maintains homeostasis in pre-cancerous tissue. The aggressiveness of colon cancer is ranked in part by the loss of patterning that has occurred within the tissue [40, 41]. Our work highlights the Erk and Wnt pathways as maintainers of patterning and therefore restrictors of dysplasia. One hypothesis is that Wnt pathway activation results in suppression of Erk during early colon tumor development. Hyperactive mutations such as oncogenic KRAS could be a selective response to Wnt activation. The result would be both the Wnt and KRAS pathways, which normally oppose one another, becoming hyperactivated together. This mutational combination, also requiring p53 suppression [42], is seen in most colon tumors, and results in highly mobile, proliferative, and adaptive clones.

The mutation frequency in the normal epithelium is far too high for DNA repair alone to explain how humans prevent expansion of cancerous cells over a lifetime [43]. How does the normal tissue prevent the takeover of hyperproliferative clones throughout life? Pathways highlighted as tumor-driving can also play a vital role in tumor prevention within normal tissue. In human cell lines, oncogenic mutations that trigger sustained Erk signaling result in extrusion carried out by normal neighbors in cell mixing experiments. This surveillance mediated by normal cells is mediated by Erk waves [18] and is an example of normal cells using a pathway described regularly as oncogenic to prevent establishment of transformed clones. Similarly, epithelial cells rely on Wnt signaling to extrude pre-cancerous cells that have acquired Wnt pathway activating mutations [44]. Our work highlights how Wnt and Erk can regulate the size and spacing of cellular compartments in the normal tissue and shows how delicately these pathways are balanced. Future work monitoring Erk and other central signaling pathways across many tumor PDOs with defined genetic backgrounds will unveil how patterning and signaling between single cells is dysregulated during tumor progression.

---

**Movie 1**

Self-organization of 2D monolayers. Time course imaging of H2B (left) and Brightfield (right) of self-organization over 120 hours. Images were acquired every 15 minutes.

**Movie 2**

Erk waves originate from apoptotic cells. Channels from left to right: H2B (purple), ErkKTR (white), Cleaved caspase 3 (green), and brightfield taken over the course of a single wave event. The red expanding circle indicated the area of high Erk activation. The green circle indicates cells that turn off Erk signaling after the initial wave passes. Images were acquired every 15 minutes for 75 minutes.

**Movie 3**

Cell Movement is instructed by Erk signaling wave. Time course imaging of a single Erk wave with or without treatment with 10µM of the EGFR inhibitor Gefitinib. Channels from left to right: H2B (purple) and ErkKTR (white). Images were acquired every 15 minutes for 105 minutes. Movie 3a is control and Movie 3b is treated.

**Movie 4a**

Erk activity is increased in non-node regions. Time course imaging of a single node. Channels from left to right: H2B (purple) and ErkKTR (white). Images were acquired every 7 minutes for 10 hours.

**Movie 4b**

Spatial heat map of Erk activity over time of a single node. Nuclear to cytoplasmic ratio of the ErkKTR was measured at the single cell level over time. A threshold for active Erk cells was then combined with spatial clustering to identify Erk-active compartments. High Erk compartments were then superimposed from each imaging timepoint to create a single heat map for regional Erk activity over time. Images were acquired every 7 minutes for 16 hours. Yellow indicates high Erk activity, blue indicates low Erk activity.

**Movie 5**

Activation of ErkKTR using Phorbol 12-myristate 13-acetate (PMA). Organoids were treated with 100nM PMA for 5 minutes prior to imaging at 20 minute intervals for 120 minutes. Channels from left to right: H2B (purple) and ErkKTR (white).

**Movie 6**

1 minute pulse of Phorbol 12-myristate 13-acetate (PMA) induces lasting node loss and transient Erk activation. Channels from left to right: H2B (purple) and ErkKTR (white). Organoids were treated with 100nM PMA for 1 minute prior to imaging. Images were acquired every 30 minutes for 48 hours.

**Movie 7**

ErkKTR is activated by Wnt deprivation. Time course imaging of a single node. Channels from left to right: H2B (purple) and ErkKTR (white). Images were acquired every 30 minutes for 23 hours. Organoids were incubated with or without recombinant Wnt3a for 24 hours prior to imaging. Movie 7a is control and Movie 7b is treated.

**S1.**
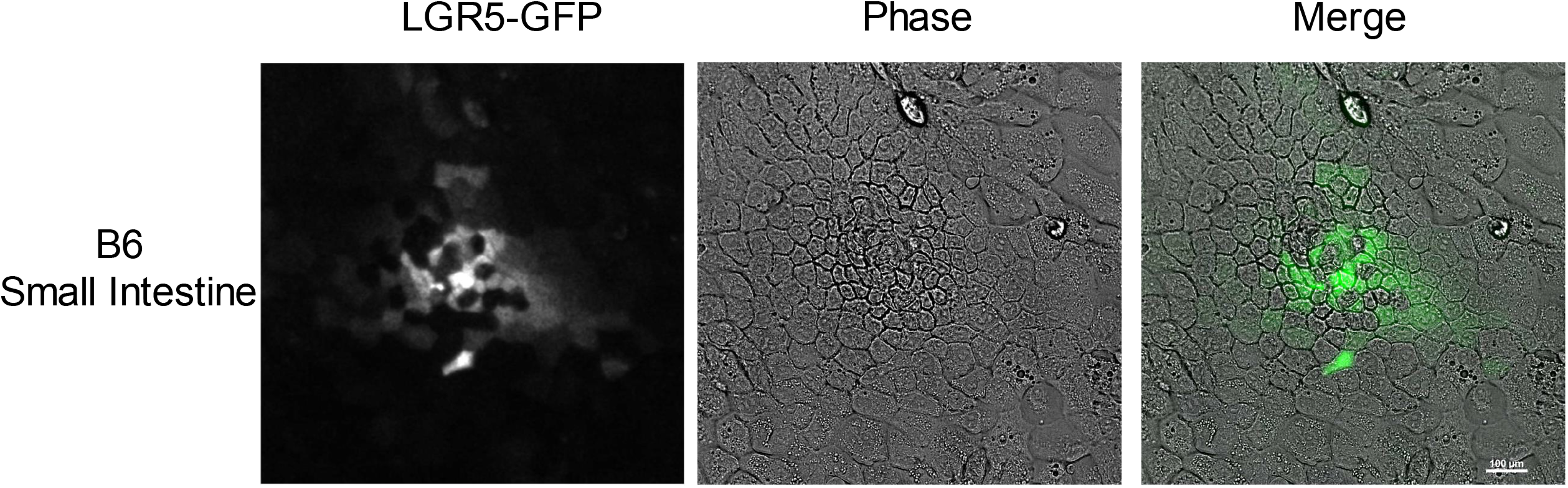
Representative images of LGR5 expressing murine small intestine organoid monolayer. Channels from left to right: LGR5-GFP (green) and phase contrast.

**S2.**
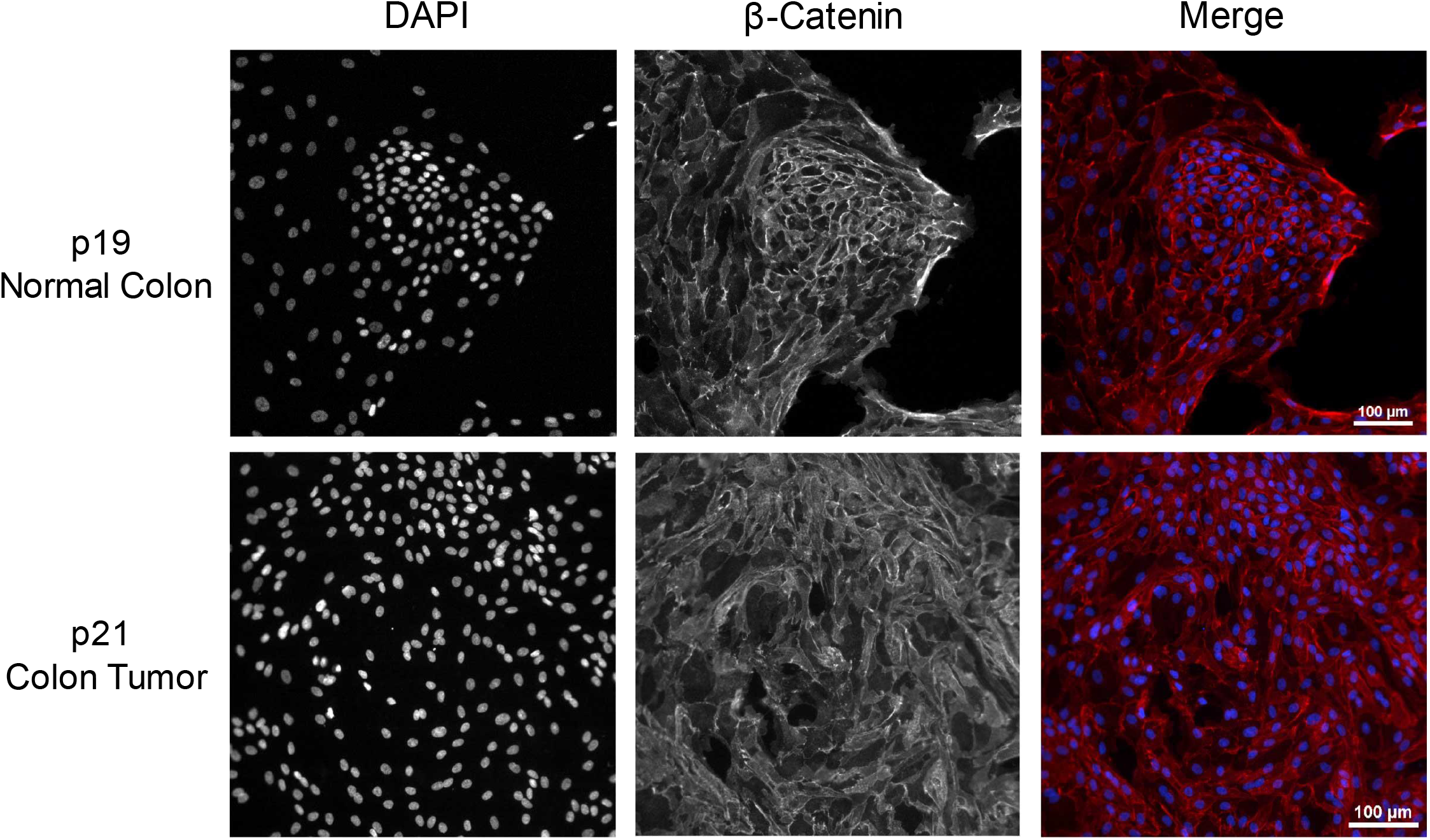
Images of organoid monolayers derived from normal (top) or tumor (bottom) derived colon tissue. Channels from left to right: DAPI (blue) and β-catenin (red).

**S3.**
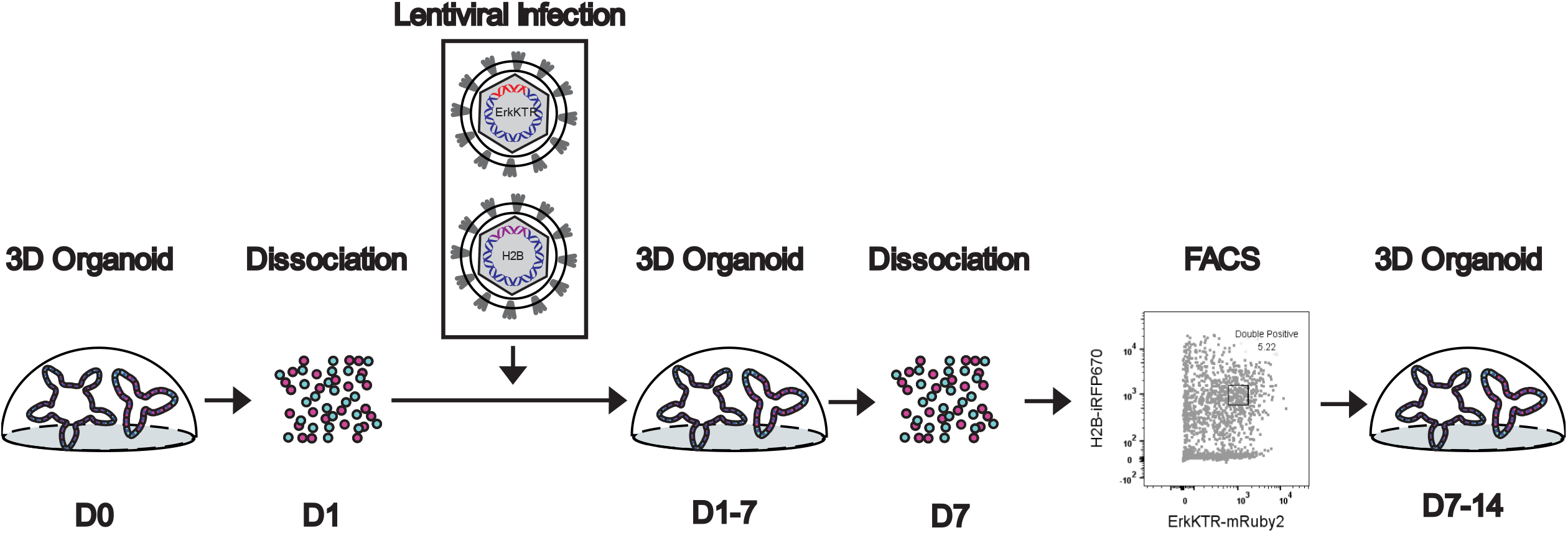
Model depicting the development of organoid monolayers expressing live cell reporters for kinase activity.

**S4.**
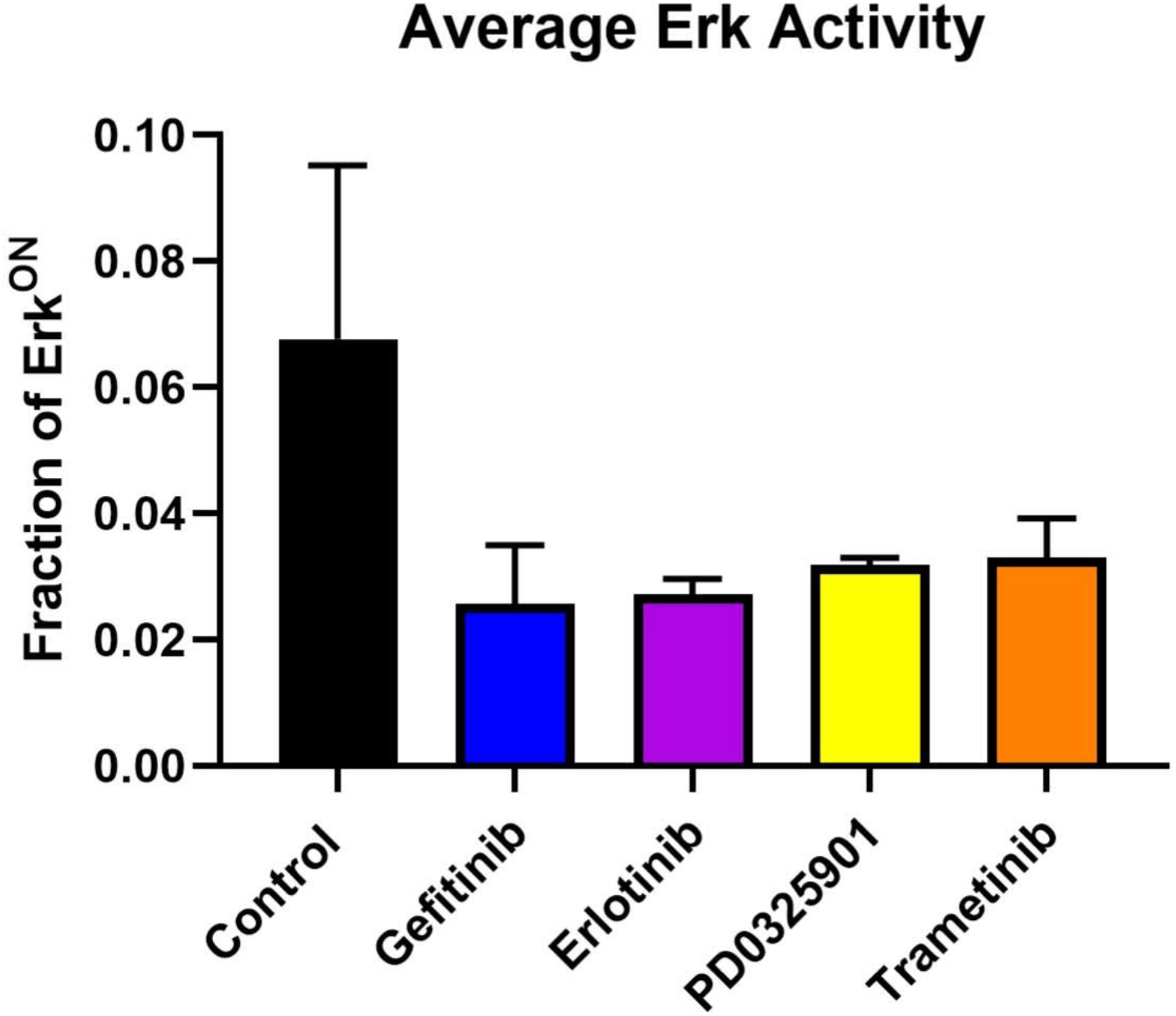
ErkKTR activity in organoid monolayers. Cells were treated with 1µM Gefitinib (EGFRi), Erlotinib (EGFRi), PD0325901 (MEKi), or Trametinib (MEKi) for 24 hours. Average Erk activity across all cells over 24 hours is shown. Error bars represent standard deviation across three independent replicates.

**S5.**
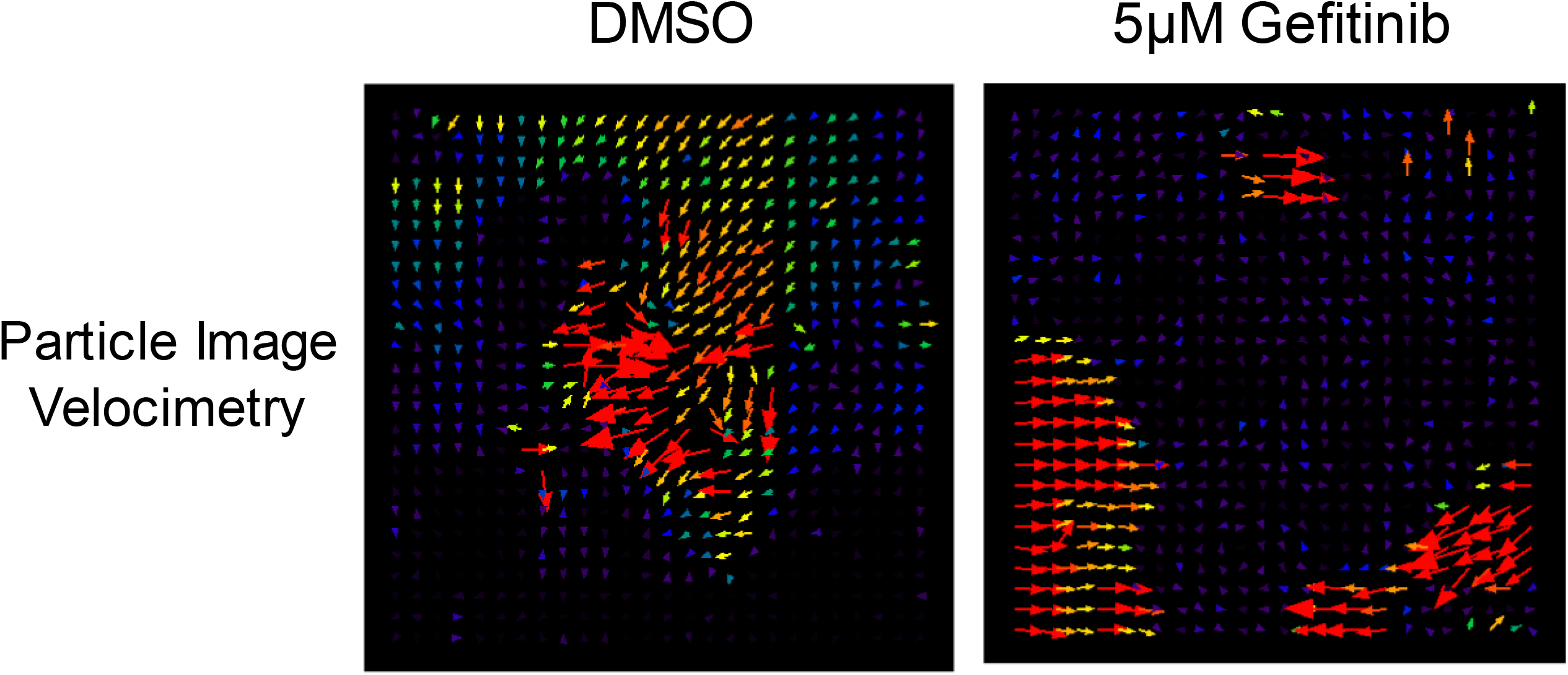
Representative images of particle image velocimetry (PIV) following cell movement after an apoptotic event. Organoids were treated with or without 5µM Gefitinib. These data correlate with movie 3. Arrows indicate direction and amplitude of movement compared to previous frame.

**S6.**
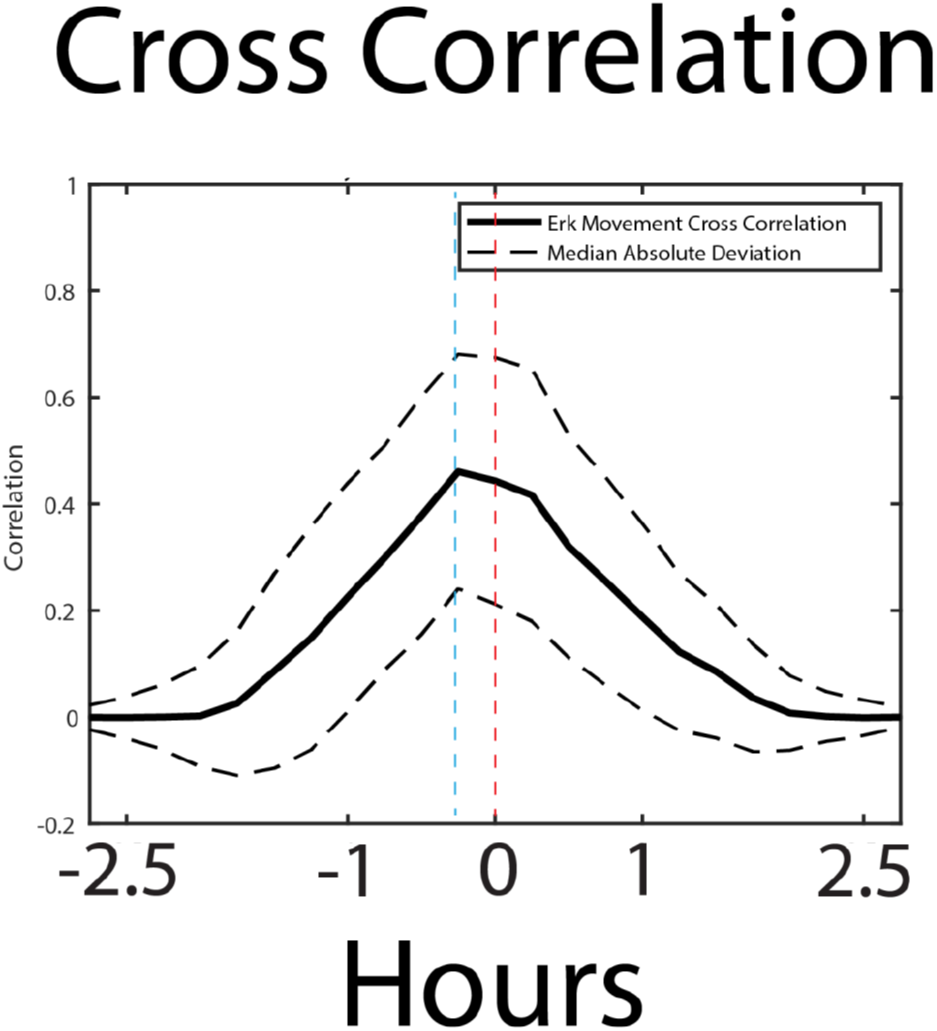
Cross correlation analysis of cell movement vs Erk activity. Correlation is shown as a solid line and the median absolute deviation is shown as dashed lines.

**S7.**
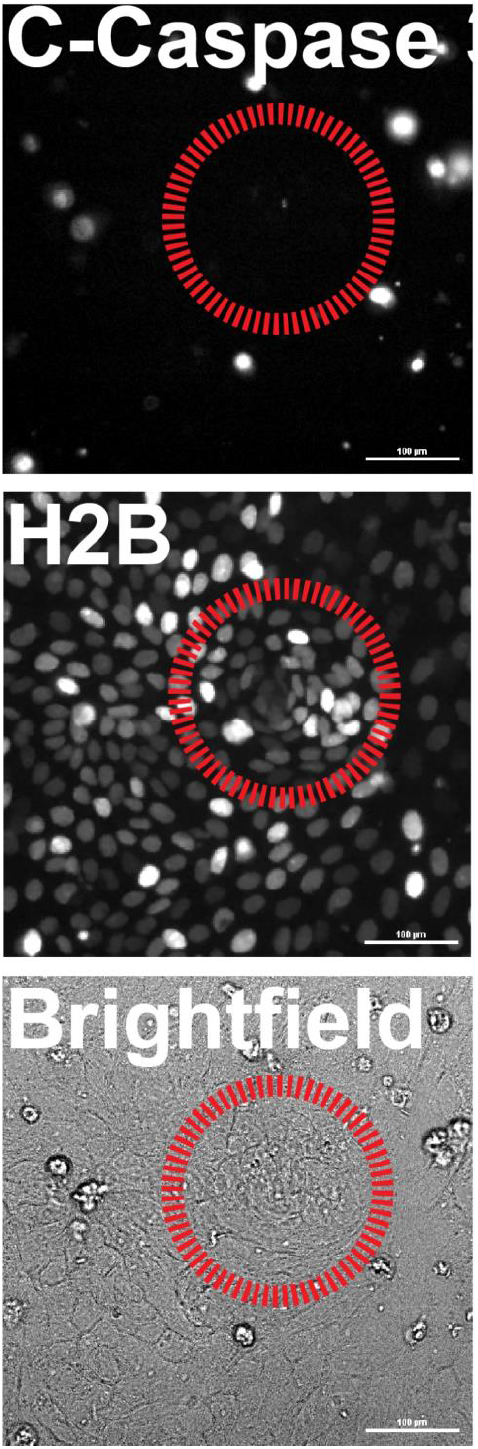
Representative images of Caspase 3 dye localization in relation to node vs non node areas. Red circle represents node area. Channels from top to bottom: cleaved caspase 3, H2B, and Brightfield. Scale bars represent 100µM.

**S8.**
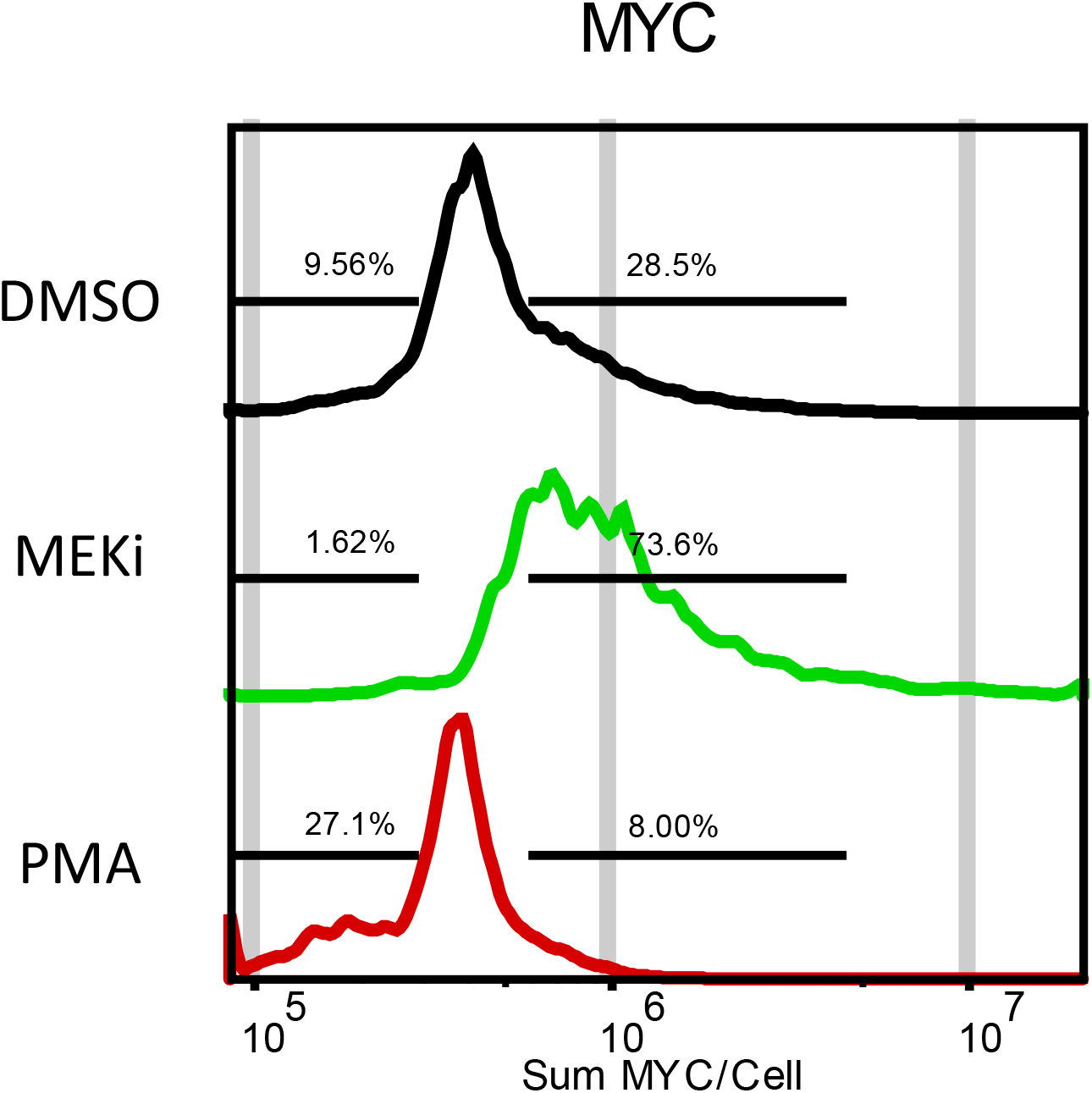
Quantification of MYC transcript levels following treatment with either 100nM PMA or 1µM PD0325901 (MEKi). Data is represented as sum intensity of the FISH probe in each cell. Gates we made against untreated control to display % population shift.

**Table 1.**
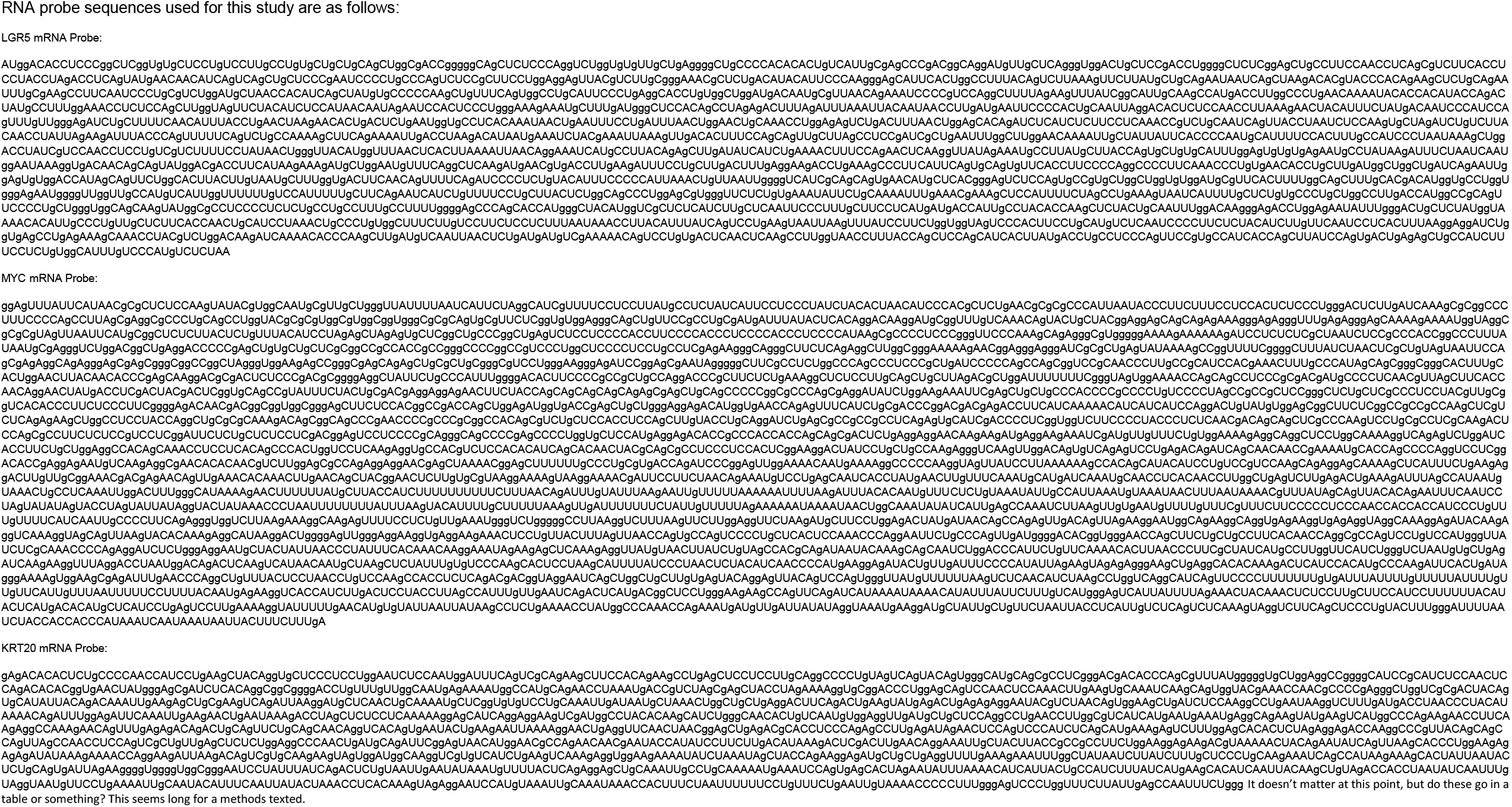
scRNA FISH custom probe sequences used for KRT20, MYC, and KRT20 are shown.

